# Disentangling effects of climate and land use on biodiversity and ecosystem services – a multi-scale experimental design

**DOI:** 10.1101/2021.03.05.434036

**Authors:** Sarah Redlich, Jie Zhang, Caryl Benjamin, Maninder Singh Dhillon, Jana Englmeier, Jörg Ewald, Ute Fricke, Cristina Ganuza, Maria Haensel, Thomas Hovestadt, Johannes Kollmann, Thomas Koellner, Carina Kübert-Flock, Harald Kunstmann, Annette Menzel, Christoph Moning, Wibke Peters, Rebekka Riebl, Thomas Rummler, Sandra Rojas Botero, Cynthia Tobisch, Johannes Uhler, Lars Uphus, Jörg Müller, Ingolf Steffan-Dewenter

## Abstract

1. Climate and land-use change are key drivers of environmental degradation in the Anthropocene, but too little is known about their interactive effects on biodiversity and ecosystem services. Long-term data on biodiversity trends are currently lacking. Furthermore, previous ecological studies have rarely considered climate and land use in a joint design, did not achieve variable independence or lost statistical power by not covering the full range of environmental gradients.
2. Here, we introduce a multi-scale space-for-time study design to disentangle effects of climate and land use on biodiversity and ecosystem services. The site selection approach coupled extensive GIS-based exploration and correlation heatmaps with a crossed and nested design covering regional, landscape and local scales. Its implementation in Bavaria (Germany) resulted in a set of study plots that maximizes the potential range and independence of environmental variables at different spatial scales.
3. Stratifying the state of Bavaria into five climate zones and three prevailing land-use types, i.e. near-natural, agriculture and urban, resulted in 60 study regions covering a mean annual temperature gradient of 5.6–9.8 °C and a spatial extent of 380×360 km. Within these regions, we nested 180 study plots located in contrasting local land-use types, i.e. forests, grasslands, arable land or settlement (local climate gradient 4.5–10 °C). This approach achieved low correlations between climate and land-use (proportional cover) at the regional and landscape scale with |*r* ≤0.33| and |*r* ≤0.29|, respectively. Furthermore, using correlation heatmaps for local plot selection reduced potentially confounding relationships between landscape composition and configuration for plots located in forests, arable land and settlements.
4. The suggested design expands upon previous research in covering a significant range of environmental gradients and including a diversity of dominant land-use types at different scales within different climatic contexts. It allows independent assessment of the relative contribution of multi-scale climate and land use on biodiversity and ecosystem services. Understanding potential interdependencies among global change drivers is essential to develop effective restoration and mitigation strategies against biodiversity decline, especially in expectation of future climatic changes. Importantly, this study also provides a baseline for long-term ecological monitoring programs.

## Introduction

Human actions are threatening the interdependent yet fragile balance of the biosphere, with far-reaching consequences for the diversity of plants (Brummitt et al., 2015) and animals (Dirzo et al., 2014). As biodiversity contributes a wealth of ecological services, cascading effects and reassembly of communities jeopardize human well-being and biosphere’s resilience against current and future disturbance (Chaplin-Kramer et al., 2019; Mori et al., 2018). Many of the services, such as food provisioning, decomposition or maintenance of soil fertility, rely on biotic interactions potentially sensitive to global change. This is especially true for regulating services provided by the highly diverse class of insects: pollination and pest regulation, both shown to strongly affect food production (Dainese et al., 2019; Duffy et al., 2017). Reported losses of insect biomass and abundances across Europe and the globe are therefore particularly worrisome (Hallmann et al., 2017; Seibold et al., 2019; Wagner, 2020). Yet the full cross-taxon magnitude of declines and the relative contributions of man-made drivers remain poorly understood.

One of the greatest threats to biodiversity is land-use change, the transformation of terrestrial ecosystems for infrastructure, human settlements and the production of crops, animals and timber (Newbold et al., 2015). Landscape simplification, urbanization, deforestation, and agricultural intensification alter environmental conditions and the availability of habitats and resources, but also the structure of entire landscapes, i.e. their composition (amount of different habitat types) and configuration (spatial arrangement and patch size of habitats). Both variables are often highly correlated (Fahrig et al., 2011) and might interact in nonlinear ways (Martin et al., 2019; Redlich et al., 2018), while attempts to disentangle them may reduce the statistical power of study designs (Fig. 1). Concurrently, land-use effects on biodiversity and ecosystem services depend on spatial scaling, the degree of specialization and movement capability of taxa and ecological processes considered (Piano et al., 2020; Wiens, 1989), with important implications for population dynamics, the diversity of fungi, plants and animals, and in consequence for ecosystem functions and services (Díaz et al., 2019; Foley et al., 2005; Newbold et al., 2015). While macroecological processes such as environmental filtering determine regional species pools, species diversity and population abundances at smaller spatial scales relate to multi-habitat use, dispersal ability, resource availability and trophic interactions. For instance, large-scale urbanization reassembles terrestrial and aquatic invertebrate communities (Piano et al., 2020), but local conversion to cropland reduces species abundances and the multitrophic functional biodiversity in agroecosystems (Provost et al., 2020) with flow-on effects for pollination, pest regulation and crop productivity (Dainese et al., 2019).

**Figure 1.**
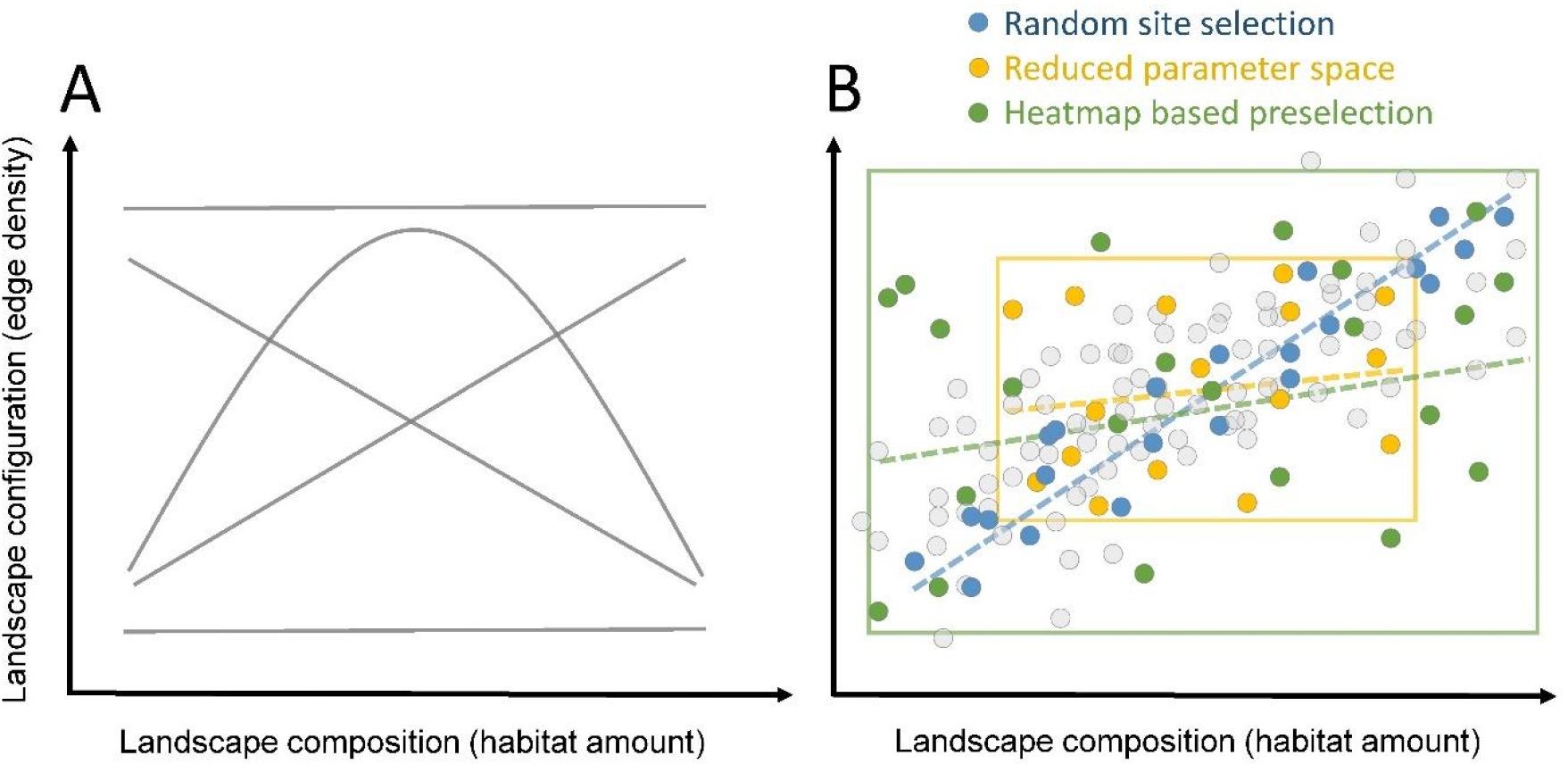
Disentangling effects of landscape composition and configuration in large-scale ecological studies. (A) Relationship between variables can be positive, negative, non-linear or independent, depending on habitat amount, habitat type and region. (B) Random selection of study plots regularly results in significant correlations between variables (blue points), while posterior exclusion of plots reduces correlations but also the covered parameter space (yellow rectangle and points). A priori knowledge of potential correlations and targeted selection of study plots using heatmaps reduces correlations and increases the parameter space (green rectangle and points). Dashed trend lines in blue, yellow and green in (B) indicate the expected change of landscape variable correlations depending on the site selection approach.

Climate is another major driver of biodiversity. Long-term data on species distributions along latitudinal and elevational climatic gradients demonstrate significant poleward and upward shifts of species’ ranges driven by global warming (Parmesan, 2006). In the future, extinction risks across all animal taxa – but particularly ectothermic organisms such as insects – may further increase with accelerating climate change (Urban, 2015; R. Warren et al., 2018). Similarly, plant community richness is likely to decrease in temperate climates, where the range of thermal tolerances in regional species pools is narrow (Harrison, 2020).

Specific land-use types may prevent climate-induced range shifts and accelerate extinctions (Fox et al., 2014; Peters et al., 2019), especially in case of less mobile specialists (Warren et al., 2001). Alternatively, (in)vertebrate communities in anthropogenic land-use types may shift towards drought- and warming-tolerant species (Williams & Newbold, 2020). Understanding the independent and combined impact of land-use and climate change on biodiversity, community composition and ecosystem services is needed to predict future changes and allow for management strategies to mitigate further losses. However, less than 10% of available studies analyse combinations of those drivers (Rillig et al., 2019). Land-use change may also feedback to the atmosphere and alter regional climate including water availability by precipitation (Dale, 1997; Laux et al., 2017; Williams & Newbold, 2020), resulting in correlated land-use and climate gradients that make it difficult to disentangle individual effects (Peters et al., 2019). Furthermore, long-term data on climate, land use and biodiversity are currently lacking, recently established monitoring schemes will not deliver sufficient data in the near future and time-series analysis may be prone to biases (Didham et al., 2020).

Here, we report on a novel protocol (Fig. 2) for a comprehensive study design that systematically combines full gradients of climate and land use at various spatial scales to investigate interacting effects on biodiversity of a wide range of taxa. This method was developed within the framework of a large-scale interdisciplinary climate research project (LandKlif, www.landklif.biozentrum.uni-wuerzburg.de). The stratified, nested design used intensive GIS-based exploration of potential study regions and a new site-selection approach based on heatmaps to reduce potential pitfalls of ecological studies on effects of land-use and climate: a) non-independence of climate and land-use variables, and correlations among land-use related composition and configuration variables; b) restrictions in gradient range or the number of spatial scales considered; c) lacking monitoring data for biodiversity and ecosystem services. The described method can be useful for similar multi-scale research programs and long-term ecosystem monitoring but will also allow for predictions of potential interactive impacts of climate and land use in a space-for-time approach.

**Figure 2.**
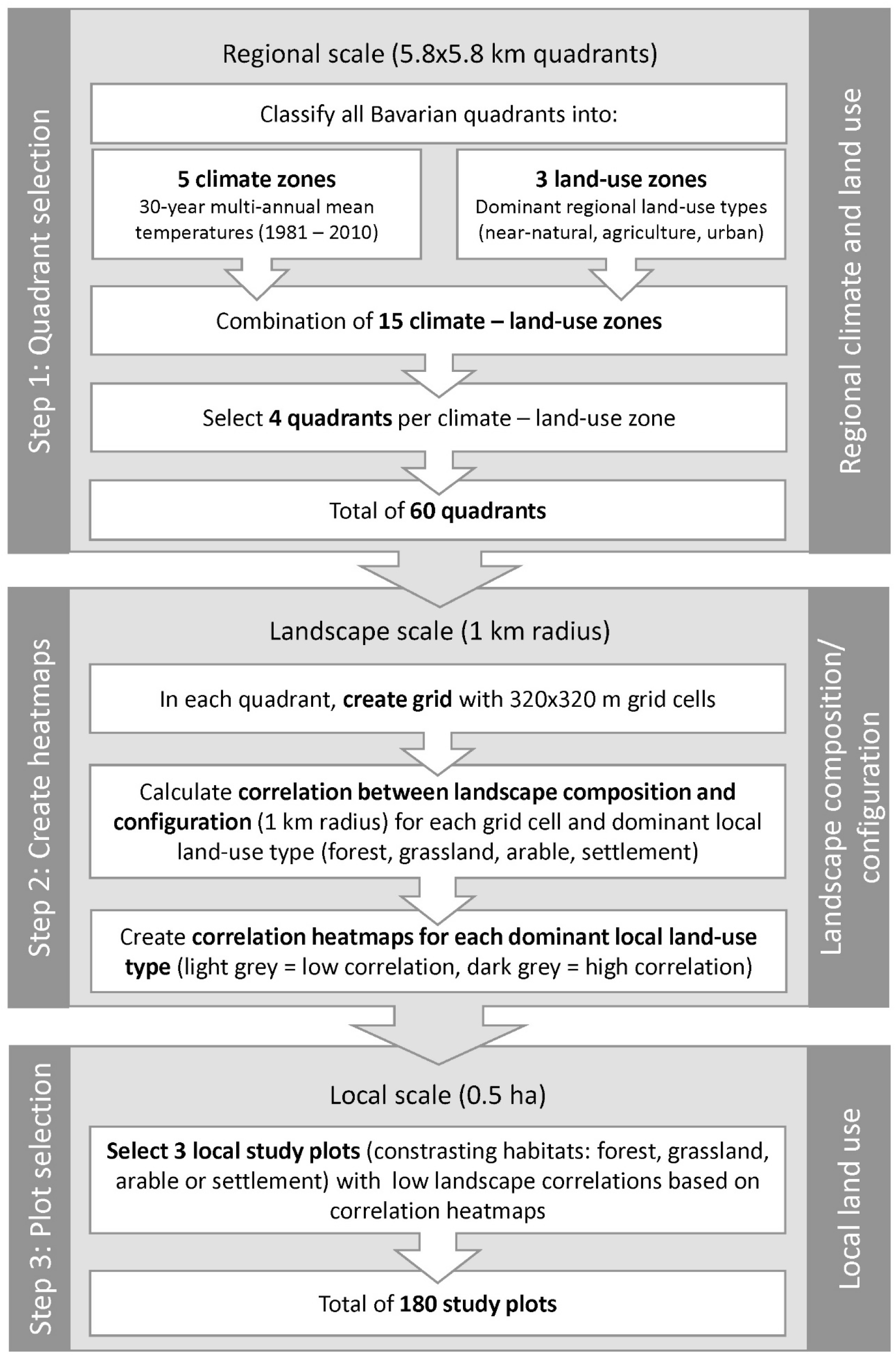
General overview of three-step plot selection process. Step 1: Selection of 60 study regions based on 15 climate – land-use combinations. Step 2: Creation of heatmaps to disentangle landscape composition and configuration variables in 1-km radius. Step 3: Based on heatmaps, selection of final 180 study plots in contrasting local land-use types.

## Material and methods

### Study area

The three-step study design (Fig. 2) was implemented in Bavaria in Southern Germany. With an area of around 70,000 km^2^ and 13 mio. inhabitants, it is the largest and second most populous state of Germany (Bayerisches Landesamt für Statistik, 2020). It covers an elevational gradient of 93–2943 m averaged at a resolution of 1 arc-second (SRTM, 2020) with mean annual temperatures (climatological reference period 1981–2010) averaged in 1-km^2^ grid cells ranging from -3.8–10.4 °C (Deutscher Wetterdienst, 2020). The land use of Bavaria is dominated by human influences, but also comprises less intensively used near- or semi-natural areas. While 7% constitute urban areas and 53% agricultural land or managed grassland, the remaining 40% are covered by (mostly managed) forests, nature protection areas and other near-natural habitats (CORINE, 2012). Bavaria’s size and heterogeneity of climate and anthropogenic influences makes it a pilot region for studying and disentangling effects of climate and land use in temperate regions and at the regional, landscape and local scale.

### Step 1 - Selection of study regions based on climate and land-use zones

At the regional scale, a stratified sampling approach ensured complete coverage of climate and land-use gradients and largely uncorrelated, orthogonal parameter combinations of both (Fig. 2). Regions were hereby defined as existing 5.8×5.8 km quadrants, which build the cells of a spatial grid covering the whole of Bavaria (‘TK25’ topographical map, scale 1:25,000). These quadrants are widely used for floristic and faunistic inventories.

To select potential climate—land-use combinations, quadrants were first classified into five climatic zones based on 30-year (1981–2010) mean air temperature data for each quadrant (Deutscher Wetterdienst, 2020). We further categorized each quadrant as one of three dominant regional land-use types based on proportional land use (CORINE, 2012): near-natural quadrants (>85% near-natural vegetation including a minimum of 50% forest), agricultural quadrants (>40% arable land and managed grassland), and urban quadrants (>14% housing, industry and traffic infrastructure). Cut-off values for land use and climate were chosen to 1) maximize climatic differences and the contrast among land-use types, with anthropogenic impact ranging from low (near-natural) to very high (urban); 2) achieve equal intervals and a similar number of quadrants within each category; and 3) obtain enough quadrants in each class to realise an even distribution and meet logistic requirements (e.g. reduce travelling time, avoid no-fly zones for UAVs where aerial assessments were planned). Based on these prerequisites, we selected four quadrants of each of the 15 climate–land-use combinations (60 study regions, Fig. 2).

### Step 2 – Create heatmaps to reduce correlations among landscape variables

Within each of the 60 study regions, we aimed to investigate the impact of local land use and interactive effects of landscape-scale land use (composition and configuration) on biodiversity and ecosystem services. The landscape-scale was hereby defined as 1-km radius around local study plots, as this scale was shown to have ecological relevance for arthropods (Bosem Baillod et al., 2017; Holzschuh et al., 2016; Thies et al., 2003). As the strength of correlations among landscape variables depends on the location of local study plots, we implemented a novel heatmap approach with a priori knowledge of potential relationships (Fig. 1). These correlation heatmaps – created for four dominant contrasting local land-use types identified within our study regions – served as systematic criterion for local study plot selection (Fig. 2).

The heatmap procedure involved the following steps: (1) Within each quadrant and starting 1 km away from the quadrant edge, we created a grid of 320 m resolution (resolution of the underlying CORINE data (2012), Fig. 4A). We calculated four landscape composition variables (proportional cover of four local land-use types: forest, grassland, arable land, settlement) and one configuration variable (edge density, i.e. length of edges between all habitat types on a per unit area, m ha^-1^) for a 1-km radius buffer around the centre of each 320×320 m grid cell (Fig. 4B). The next steps, here exemplified for forest, were repeated for each local land-use type. (2) We selected all grid cells (Fig. 4C) with a proportional forest cover of >20% (to accommodate a 0.5-ha study plot and a 3×30 m experimental area) and >5% forest in the surrounding 1-km radius buffer (to ensure a minimum amount of forest was present in the surrounding landscape). (3) Of these forest grid cells and associated landscape buffers, we randomly chose one in each of the 60 study quadrants - if existent (quadrants without forest grids were excluded) - and calculated the overall Pearson’s *r* correlation coefficient between the surrounding landscape composition (here forest cover) and configuration (edge density) based on the random plot selection. (4) This random selection and calculation was repeated 10,000 times. (5) For each forest grid-cell *i* we then calculated the average Pearson’s 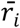 coefficient across all the random combinations of points in which this cell was included:

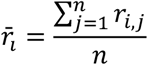

Where *r*_*i,j*_ is the *j*^th^ Pearson’s *r* coefficient resulting from random selection of that specific forest dominated grid cell *i*, and *n* is the number of times that grid cell *i* was included in one of the 10,000 random selections of points. (6) In a last step and considering all forest grid cells in our 60 quadrants, we used natural breaks (Jenks natural breaks algorithm implemented in ArcMap v10.4) to classify the range of mean correlations into three categories to create the correlation heatmap for the local land-use type forest (Fig. 4C). By repeating the steps described in (2–6) for all land-use types (forest, grassland, arable land, settlement), we derived a set of four heatmaps for each of the 60 quadrants. During the local plot selection process (Step 3), these heatmaps helped to reduce correlations of landscape composition and configuration around plots with specific land-use types (e.g. only forest plots), but also across all study plots.

### Step 3 - Selection of local study plots

Within each quadrant, we aimed to establish local study plots of 0.5 ha size within contrasting land-use types (Fig. 2). Although four local, dominant land-use types had been identified during the heatmap process (forest, grassland, arable land or settlement), not all were present in each quadrant. Therefore, we focused on three out of four land-use types per quadrant by considering availability (if only three types present) or regional dominance (three types with highest proportional cover) and contrast (whenever proportional cover of two land-use types was similar). We then used the respective heatmaps to preferentially place study plots in grid cells that had a low predicted correlation values for the specific land-use type.

Additional decision rules for plot selection included landowner permission, >2 km between plots, >50 m away from roads, water bodies and other land-use types, protection from vandalism and good accessibility. Nested within our large-scale factorial design, the resulting 180 plots allowed us to assess the influence of local land use on biodiversity and ecosystem services, while minimizing correlations between landscape composition and configuration.

### Assessing efficiency of study design

We assessed the efficiency of our stratified selection and heatmap approach by a) region (5.8×5.8 km): calculating Pearson’s *r* correlation coefficients between climate and the proportion of our regional dominant land-use types near-natural, agriculture and urban; b) landscape (1-km radius): assessing relationships between climate and the proportion of our dominant local land-use types forest, grassland, arable land and settlement. We also visually compared final correlations between landscape composition and configuration with potential correlations based on 10,000 random selections.

The proportion of land-use types (region, landscape) and landscape composition and configuration variables were calculated in ArcGIS pro v2.2.0 and ArcMap v10.4 using CORINE data (2012). Climate data for regions and landscapes (mean air temperatures and associated precipitation amounts) were calculated using Esri ASCII grid raster files with 1×1km resolution (Deutscher Wetterdienst, 2020) by averaging pixel values within each 5.8×5.8 km quadrant and 1-km buffer around selected study plots, respectively. All Pearson’s *r* coefficients calculated in R v4.0.2.

## Results

### Implementation of the experimental design

Our design and selection process (Fig. 2) allowed us to minimize the potential correlations between climate, land use and landscape metrics at multiple scales and resulted in an approximately even distribution of 60 study regions (quadrants) across Bavaria (Fig. 3).

**Figure 3.**
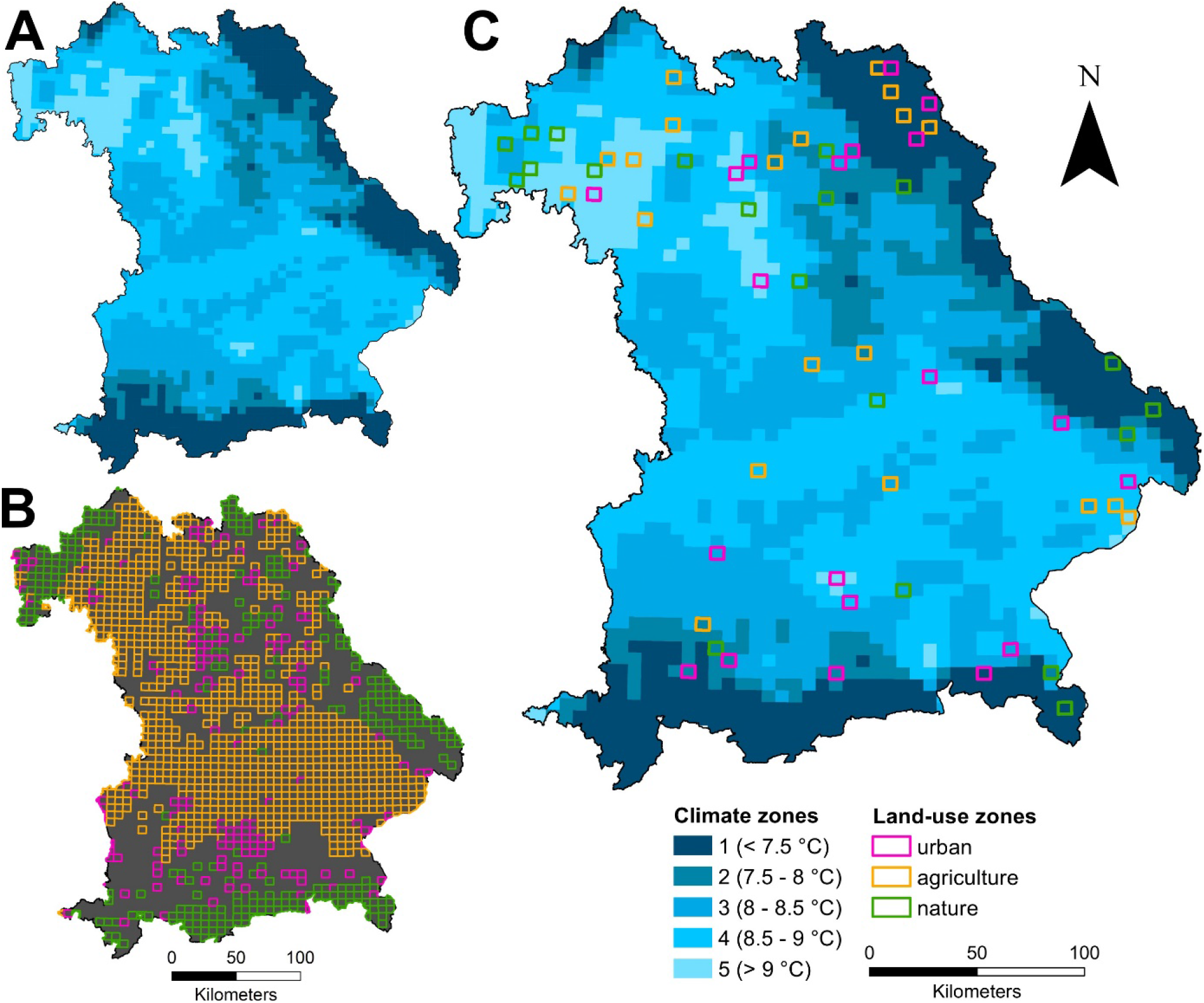
Implementation of a full-factorial, stratified design crossing regional climate and land use in Bavaria, Southern Germany. Climate zones (A) were based on 30-year (1981– 2010) mean air temperatures in each quadrant (1 (cold) to 5 (warm)). For land use (B), we distinguished between near-natural quadrants (>85% natural vegetation including a minimum of 50% forest), agricultural quadrants (>40% arable land and managed grassland) and urban quadrants (>14% housing, industry and traffic infrastructure). The final 60 study regions (C) covered 15 climate–land use combinations with four replicates each.

**Figure 4.**
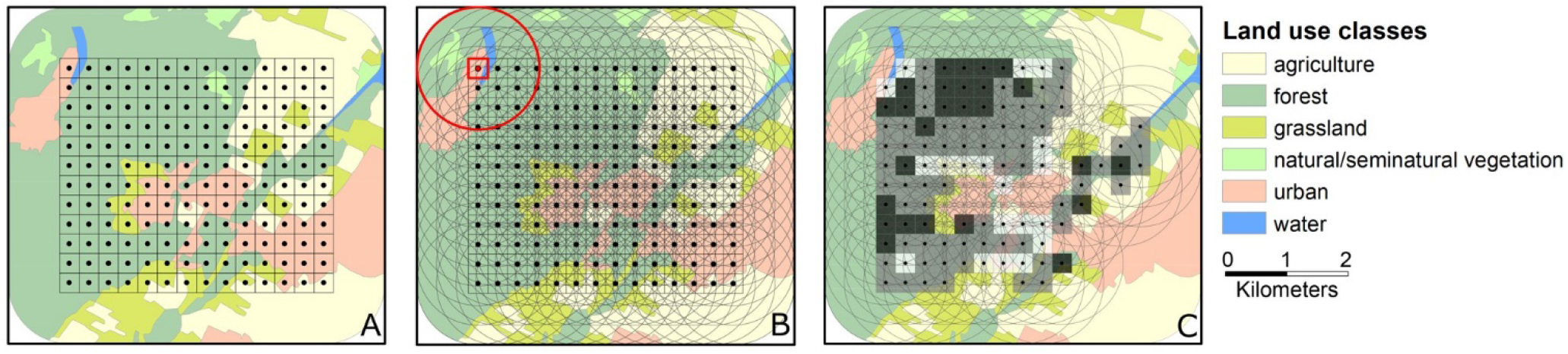
Process of deriving correlation heatmaps for each dominant land-use type to guide the selection of local study plots. Colours of polygons represent different land-use types. (A) Create a fishnet of 320 m resolution inside each of 60 study quadrants. (B) Calculate landscape composition and configuration within a 1-km radius around centre of each 320×320 m grid cell. (C) Select grid cells dominated by the respective land-use type (here forest, dark green) and create land-use specific heatmaps of mean correlations between landscape composition and configuration based on 10,000 random selections of grid cells across all quadrants. Shades of grey in heatmaps indicate levels of the predicted degree of correlation (light = high correlation, dark = low correlation) if the respective grid was chosen.

These regions covered a climate gradient of 5.6–9.8 °C (8.2 ± 0.8 °C, mean ± SD) and 614– 1820 mm of annual precipitation amounts (939 ± 263 mm). Across all quadrants, the cover of our dominant regional land–use types (i.e. landscape composition) ranged from 0.8 to 97.1% (40 ± 27.7%) for near-natural land use, 0.3–91.0% (44.7 ± 24.9%) for agriculture, and 0– 97.2% (14.7 ± 21.1%) for urban areas. Regional mean temperatures (Fig. 5A–C) and precipitation (|*r*<0.3|, Appendix Fig. S1A–C) showed low correlations with regional land use (proportion of near-natural, agriculture and urban habitat).

**Figure 5.**
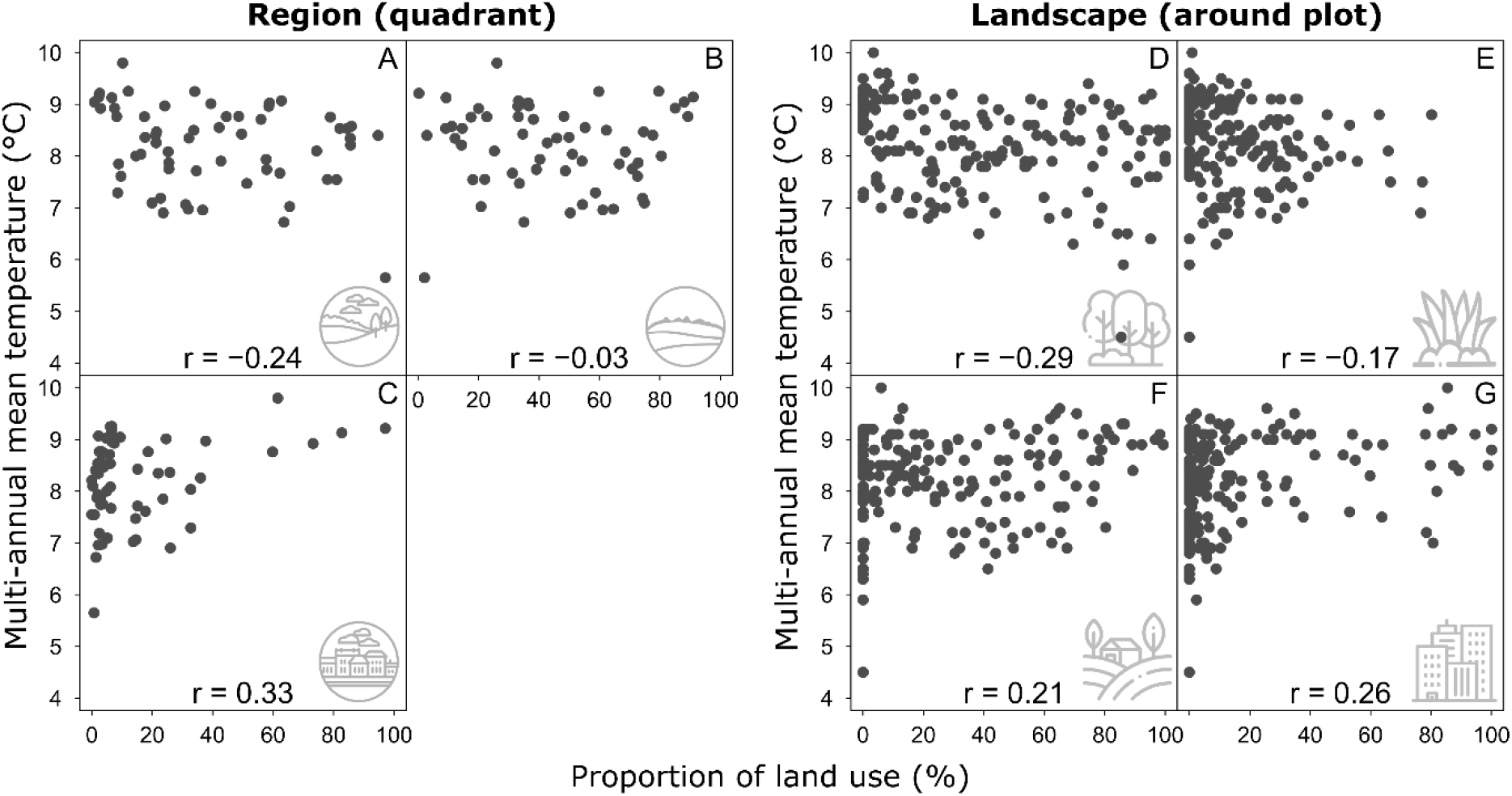
Relationships between 30-year mean temperatures (1981–2010) and proportional land cover (composition) for the regional land-use types near-natural (A), agriculture (B) and urban (C), and for the landscape-scale land-use types forest (D), grassland (E), arable land (F) and settlement (G). Pearson’s *r* coefficients based on 60 study regions (5.8×5.8 km quadrants, A–C) and 179 (out of expected 180) study plots (1-km radius around local study plots, D–G).

For each study region, the heatmap procedure yielded four heatmaps for the local land-use types forest, grassland, arable land and settlement, which were used to identify potential study plots within dominant local land-use types (Fig. 6B–D). After ground-truthing of sites and gaining permission of landowners, three final plots were chosen per quadrant (Fig. 6E), yielding 179 out of 180 expected study plots (Fig. 6A). One study plot was discarded as landowner permission was denied. Forest (*n* = 55) was the most selected local land-use type, followed by grassland (*n* = 46), arable land (*n* = 43) and settlement (*n* = 35).

**Figure 6.**
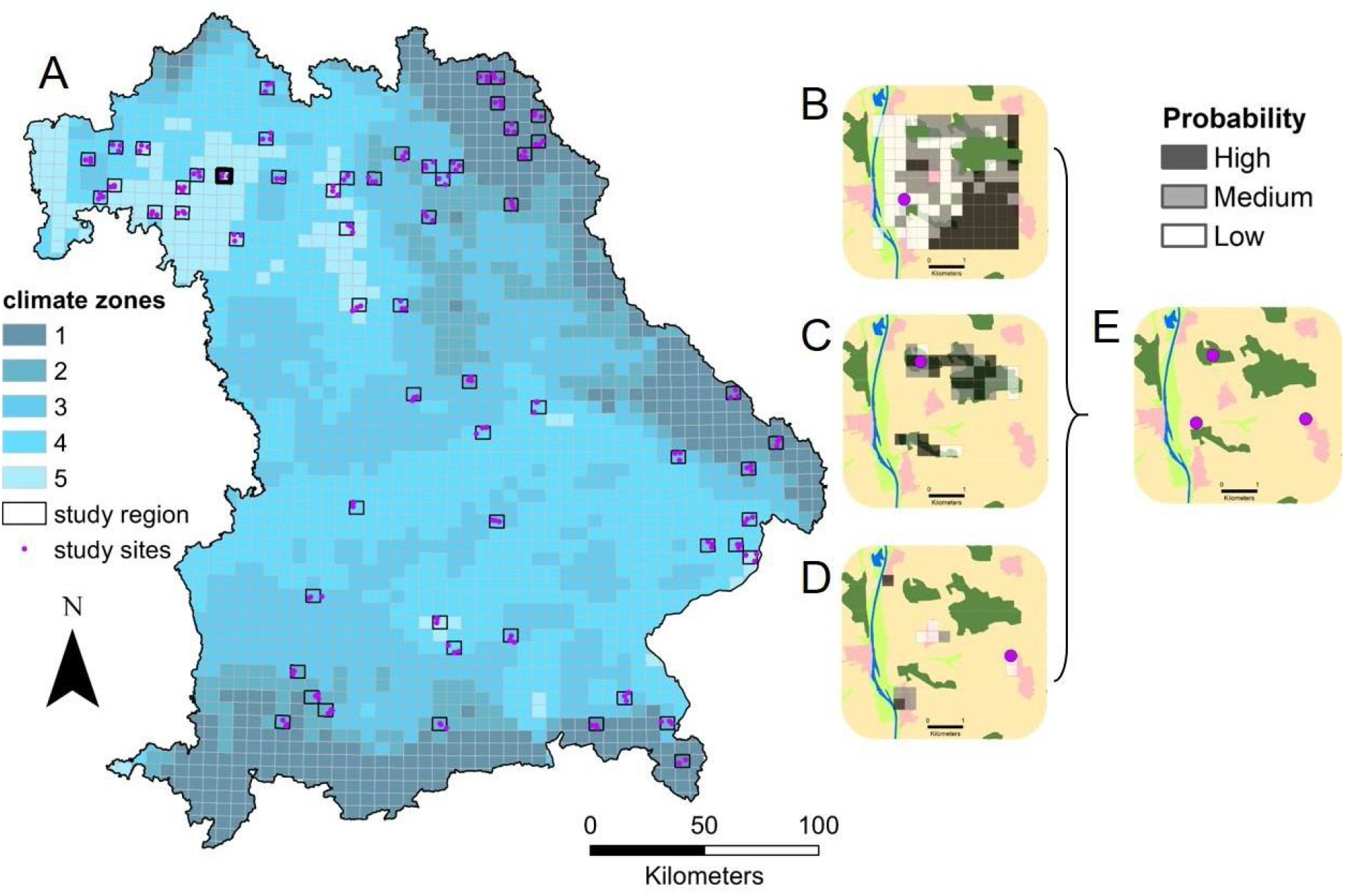
: Map of all 179 (out of expected 180) study plots in 60 study regions (A) and example of heatmaps for three dominant local land-use types (arable land (B), forest (C), and settlement (D)) used for the final selection of study plots (E). Shades of grey in heatmaps indicate levels of the predicted degree of correlation (light = high correlation, dark = low correlation) if the respective grid was chosen.

At the landscape-scale in 1-km radius around study plots, mean temperatures across our 179 study plots ranged from 4.5–10 °C (8.2 ± 0.8 °C), with annual precipitation amounts of 590–2893 mm (933 ± 279 mm). Landscape composition gradients across all plots stretched from 0–100% for forest (37.9 ± 32.3%), 0–80.2 % for grassland (15.7 ± 17.1%), 0–99.4% for arable land (28.7 ± 29.2%) and 0–100% for settlement (16.1 ± 25.8%). Edge density across all study plots was 0–66.0 m ha^-1^ (28.1 ± 13.8 m ha^-1^). Correlations of landscape-scale temperature with composition variables (Fig. 4D–G) and edge density were low (Pearson’s *r* = - 0.17).

Compared to potential correlations based on random selection of study plots, the heatmap approach resulted in lower correlations between landscape composition and configuration (in 1-km radius around study plots) for plots located in forest, arable land and settlements (blue line, Fig. 7A, C, D). Only for grassland, the final correlation was positive and higher than predicted (blue line, Fig. 7B). Taking all study plots independent of the local land-use type into account, this pattern was even stronger, with correlations between the proportion of habitats and edge density being very low for forest (Pearson’s *r* = -0.31), arable land (*r* = 0.09) and settlement (*r* = -0.08), yet high for grassland (*r* = 0.51) (red line, Fig 7, Fig. S2). Correlations among composition variables ranged from *r* = -0.13 (settlement and grassland) to -0.55 (arable land and forest).

**Figure 7.**
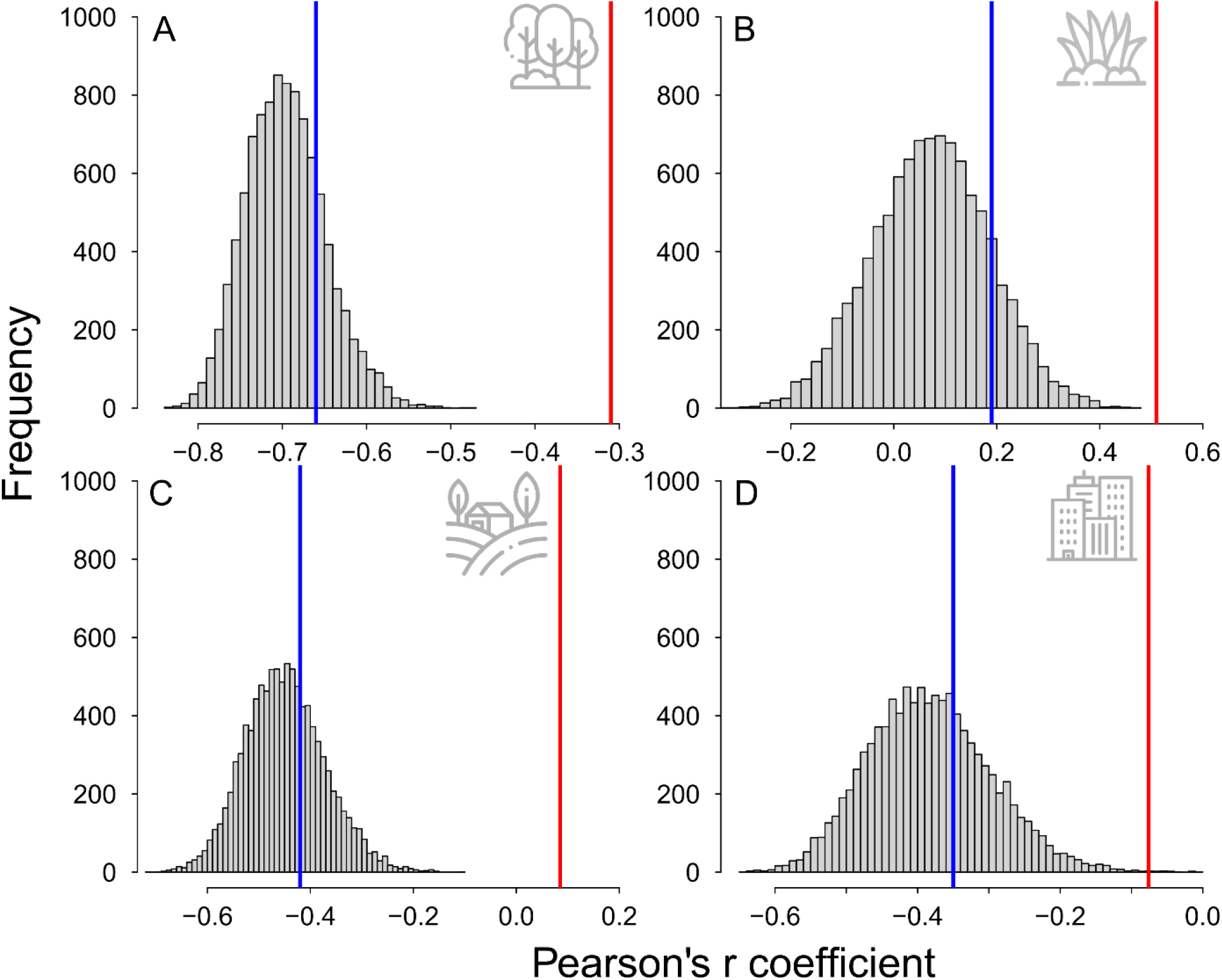
Potential and actual Pearson’s correlations between landscape composition (proportional cover of land-use types) and configuration (edge density) in 1-km radius around study plots. Compared to the histograms of potential correlations resulting from 10,000 random selections of grid cells (i.e. potential study plots, cf. ‘heatmap procedure’), blue lines show reduced actual correlations based on subsets of plots located in the land-use types forest (*n* = 55, A), arable land (*n* = 43, C) and settlement (*n* = 35, D), yet higher correlations for plots located in grassland (*n* = 46, B). Red lines show correlations for land-use specific actual correlations across all selected study plots (*n* =179).

This multi-scale GIS-supported study design is suited to disentangle climate and land-use effects on general and functional biodiversity and plant- or animal-based ecosystem services, as done within this project using a range of observational, empirical, modelling and survey data collected on different spatial scales in 2019 and 2020 (Table 1).

**Table 1:** Example for assessments of biodiversity, ecosystem services and socio-economic/management information in the LandKlif project. Observational and empirical data was collected on up to 179 study plots in 2019 and 2020 and complemented with modelling approaches and stakeholder surveys. Extended categorization of ecosystem services based on TEEB (2010) and Rabe et al. (2016).

|  | Group/Service | Detail | Scale |
| --- | --- | --- | --- |
| Biodiversity | Plants | Plant/Pollen diversity and phenology | Plot |
|  | Microbes | Soil/decomposer microbial diversity | Plot |
|  | Arthropods | Total Biomass and richness of flying and crawling arthropods; functional abundance and richness of arthropod decomposers, pollinators, trap-nesting Hymenoptera, pests and predators | Plot |
|  | Vertebrates | Diversity and density of game | Plot/Bavaria |
|  | Ecosystems | Landscape diversity, composition, configuration | Landscape/Region |
| Ecosystem services | Decomposition | Decomposition of deadwood, carrion and dung | Plot |
|  | Pest regulation | Predation and parasitism rate, herbivory | Plot |
|  | Pollination | Seed set and pollination services | Plot/Region |
|  | Productivity | Crop biomass and yield; vegetation biomass; Normalized Difference Vegetation Index (NDVI); flower resource availability | Plot/Region |
|  | Soil fertility | Soil organic carbon and nutrient content | Plot |
|  | Soil erosion prevention | Erosion | Region |
|  | Carbon sequestration | Soil Organic Carbon | Bavaria |
|  | Microclimate regulation | Temperature | Plot |
|  | Flood control | Prevention of floods | Region |
|  | Water quality regulation | Nitrogen and phosphorus retention | Region |
| Other | Stakeholder preferences | Preferences for ecosystem services | Region |
|  | Stakeholder perceptions | Climate change perceptions | Region |
|  | Landowner management | Management of land and crop fields used for experiments | Plot |

## Discussion

Studies assessing the combined effects of land use and climate on biodiversity and ecosystem services commonly struggle with non-independence of climate and land-use variables, restrictions in gradient range or scale and insufficient long-term data sets. Here, we present the protocol for a large-scale experimental design that aims to overcome these issues. While our basic design follows the selection principles for multi-scale landscape studies outlined in previous papers (Fahrig et al., 2011; Gillespie et al., 2017; Pasher et al., 2013), the use of a novel, automated heatmap approach and the inclusion of independent climatic gradients sets this design apart, both as baseline and space-for-time study.

First, the crossed and nested design at the regional scale resulted in relatively weak correlations between climate and land use (proportional cover of forest, near-natural and urban area). The design also decoupled regional climate and land-use effects from the influence of small-scale land use due to the selection of three out of four dominant local land-use types (forest, grassland, arable land or settlements) within our 60 study regions.

Regarding landscape composition and configuration in a 1-km radius around study plots, the heatmap approach lowered correlations compared to average potential correlations for specific local land-use types (blue lines, Fig. 7), but these benefits were not that substantial in absolute terms (i.e. correlations for selected plots quite close to peak of distribution for random selection). However, there are three points to consider: 1) these actual correlations were based on a subset of plots (specific local land-use types), and were much lower for forest, arable land and settlement if calculated across all study plots (red lines, Fig. 7), which is the gradient range primarily used for analysis in our project; 2) reducing landscape correlations may be difficult for land-use types such as forest, where patches generally occur clustered, causing higher negative correlations with edge density than for settlements or arable land. For grassland, correlations seem the be generally low, yet increased during the selection process, possibly due to inherent correlations among land-use types and non-linear relationships between grassland amount and edge density in the landscape; 3) in our project, complex private ownership structures, logistic and other constraints (e.g. transportation costs, time constraints, accessibility, permissions) prevented us from selecting combinations of study plots closer to *r*=0. Our method is situated halfway between two extremes: the blind selection of study plots that may inherently cause strong landscape correlations or requires the reduction in parameter space (see Fig. 1) and choosing the best available random selection of plots during the process of creating heatmaps. Accordingly, the chance of moving towards low landscape correlations ultimately depends on the gradient range and land-use type considered and methodological, logistical and ownership constraints that may be lower in other studies.

Second, we increased the coverage of spatial scales and land-use types, thereby maximizing the number of explanatory factors that can be analysed in parallel. Concurrently, our method of ‘a priori’ employing long-term climate data and extensive GIS-based exploration of potential study plots enabled us to cover independent, large climatic and land-use gradients. For landscape composition and configuration of the full set of 179 final study plots, our data highlights the natural, unimodal relationship between these variables, which is most pronounced for forest cover and grows weaker from grassland to arable land and settlement, with peaks between 40–60% land cover (Appendix S2). This implies that studies covering narrow landscape gradients between 0–50% or 50–100% may observe contrasting positive or negative correlations between these landscape variables, respectively, while studies focussed on intermediate landscape gradients are most likely to reduce the correlation between variables and differentiate between individual effects, which may be impossible at the extreme ends of the spectrum.

Finally, our extensive on-field assessments within this experimental framework will fill existing knowledge gaps about biodiversity trends across taxa, relationships between above- and belowground arthropods and the microbial diversity of decomposer communities. We can also assess potential trade-offs among ecosystem service provisioning and current and predicted interactive effects of climate and land use on biodiversity-ecosystem functioning relationships. In this context, the implemented space-for-time approach has crucial advantages over time series. Recently established long-term biodiversity monitoring schemes will not yield meaningful results before several decades, which may be too late considering the current speed of global change. Furthermore, long-term climatic change often goes hand in hand with land-use change, making it difficult to disentangle individual effects (Dale, 1997). In addition, issues such as shifting baselines or phenologies, bias in site selection and detection may cause misleading results in time series analysis (Didham et al., 2020). Other methods, such as large-scale, manipulative climate–land-use experiments following the idea of BACI designs (Before-After-Control-Impact studies, Christie et al., 2019) are highly interesting but almost impossible to implement.

Space-for-time approaches also have limitations. For instance, other drivers of biodiversity, such as anthropogenic pressure or altered biotic interactions, may mask the response to climate, especially if only small spatial scales (a few kilometres or less) with small climatic differences are considered (Blois et al., 2013). In contrast, data obtained from spatial observations was shown to overestimate phenology responses to temperature compared to long-term phenological data (Jochner et al., 2013). Still, space-for-time substitutions based on the largest possible climatic gradient is a useful and fast alternative to gain important, policy-relevant insights into the interactive effects of climate and land-use change on biodiversity and ecosystem services. By utilizing the full parameter space of the climatic and landscape variables assessed here (Fig. 1), we enhanced the validity of space-for-time substitutions related to climate change (Blois et al., 2013). We further reduced the chance of observing misleading findings in cases where non-monotonic relationships cause contradictory relationships between environmental variables and biodiversity if only a narrow variable range is used (Eigenbrod et al., 2011).

## Conclusions

Our multi-scale study protocol expands on previous designs which addressed local gradients in climate and land use (Peters et al., 2019) or gradients in landscape structure in multiple regions (Gillespie et al., 2017; Holzschuh et al., 2016). It allows to evaluate scale-dependent and interactive effects of current climate and land-use gradients on biodiversity and ecosystem services, and to predict long-term responses to climate change. Furthermore, it provides valuable baseline data to assess the effectiveness of future restoration measures at local, landscape and regional scales. We believe that this approach of an objective, multi-scale site selection across large regions deserves consideration in the implementation of national and European long-term ecosystem monitoring schemes.

## Acknowledgements

Icons used in graphs made by Freepik (http://www.flaticon.com/). CORINE Land Cover (CLC) provided by the European Union, European Environment Agency (EEA) under the framework of the Copernicus programme. Climate data provided by Deutscher Wetterdienst (DWD). This study was conducted within the framework of the joint project *Landklif* (https://www.landklif.biozentrum.uni-wuerzburg.de/) funded by the Bavarian Ministry of Science and the Arts via the Bavarian Climate Research Network (bayklif).

## Authors’ contributions

SR, JZ, JM, TH and ISD conceived the ideas and designed the methodology; JZ and CKF collected the data; SR and JZ analysed the data; SR, JZ and ISD led the writing of the manuscript. All authors contributed critically to the drafts and gave final approval for publication.

## Data Availability

Data available from the Dryad Digital Repository http://XXX (Redlich et al 2021).

## References

Bayerisches Landesamt für Statistik. (2020). Statistics. https://www.statistik.bayern.de/

Blois, J. L., Williams, J. W., Fitzpatrick, M. C., Jackson, S. T., & Ferrier, S. (2013). Space can substitute for time in predicting climate-change effects on biodiversity. Proceedings of the National Academy of Sciences, 110(23), 9374–9379. https://doi.org/10.1073/pnas.1220228110

Bosem Baillod, A., Tscharntke, T., Clough, Y., & Batáry, P. (2017). Landscape-scale interactions of spatial and temporal cropland heterogeneity drive biological control of cereal aphids. Journal of Applied Ecology, 1804–1813. https://doi.org/10.1111/1365-2664.12910

Brummitt, N. A., Bachman, S. P., Griffiths-Lee, J., Lutz, M., Moat, J. F., Farjon, A., Donaldson, J. S., Hilton-Taylor, C., Meagher, T. R., Albuquerque, S., Aletrari, E., Andrews, A. K., Atchison, G., Baloch, E., Barlozzini, B., Brunazzi, A., Carretero, J., Celesti, M., Chadburn, H., … Lughadha, E. M. N. (2015). Green Plants in the Red: A Baseline Global Assessment for the IUCN Sampled Red List Index for Plants. PLOS ONE, 10(8), e0135152. https://doi.org/10.1371/journal.pone.0135152

Chaplin-Kramer, R., Sharp, R. P., Weil, C., Bennett, E. M., Pascual, U., Arkema, K. K., Brauman, K. A., Bryant, B. P., Guerry, A. D., Haddad, N. M., Hamann, M., Hamel, P., Johnson, J. A., Mandle, L., Pereira, H. M., Polasky, S., Ruckelshaus, M., Shaw, M. R., Silver, J. M., … Daily, G. C. (2019). Global modeling of nature’s contributions to people. Science, 366(6462), 255–258. https://doi.org/10.1126/science.aaw3372

Christie, A. P., Amano, T., Martin, P. A., Shackelford, G. E., Simmons, B. I., & Sutherland, W. J. (2019). Simple study designs in ecology produce inaccurate estimates of biodiversity responses. Journal of Applied Ecology, 56(12), 2742–2754. https://doi.org/10.1111/1365-2664.13499

CORINE. (2012). Copernicus Land Monitoring Service 2012, European Environment Agency. https://land.copernicus.eu/pan-european/corine-land-cover/clc-2012

Dainese, M., Martin, E. A., Aizen, M. A., Albrecht, M., Bartomeus, I., Bommarco, R., Carvalheiro, L. G., Chaplin-Kramer, R., Gagic, V., Garibaldi, L. A., Ghazoul, J., Grab, H., Jonsson, M., Karp, D. S., Kennedy, C. M., Kleijn, D., Kremen, C., Landis, D. A., Letourneau, D. K., … Steffan-Dewenter, I. (2019). A global synthesis reveals biodiversity-mediated benefits for crop production. Science Advances, 5(10), eaax0121. https://doi.org/10.1126/sciadv.aax0121

Dale, V. H. (1997). The Relationship Between Land-Use Change and Climate Change. Ecological Applications, 7(3), 753–769. https://doi.org/10.1890/1051-0761(1997)007[0753:TRBLUC]2.0.CO;2

Deutscher Wetterdienst. (2020). DWD Climate Data Center (CDC): Multi-annual means of grids of precipitation and air temperature (2m) over Germany from 1981-2010, version v1.0. https://opendata.dwd.de

Díaz, S., Settele, J., Brondízio, E., Ngo, H., Guèze, M., Agard, J., Arneth, A., Balvanera, P., Brauman, K., Butchart, S., Chan, K., Garibaldi, L., Ichii, K., Liu, J., Subrmanian, S., Midgley, G., Miloslavich, P., Molnár, Z., Obura, D., … Zayas, C. (2019). Summary for policymakers of the global assessment report on biodiversity and ecosystem services of the Intergovernmental Science-Policy Platform on Biodiversity and Ecosystem Services. Secretariat of the Intergovernmental Science-Policy Platform on Biodiversity and Ecosystem Services.

Didham, R. K., Basset, Y., Collins, C. M., Leather, S. R., Littlewood, N. A., Menz, M. H. M., Müller, J., Packer, L., Saunders, M. E., Schönrogge, K., Stewart, A. J. A., Yanoviak, S. P., & Hassall, C. (2020). Interpreting insect declines: Seven challenges and a way forward. Insect Conservation and Diversity, 13(2), 103–114. https://doi.org/10.1111/icad.12408

Dirzo, R., Young, H. S., Galetti, M., Ceballos, G., Isaac, N. J. B., & Collen, B. (2014). Defaunation in the Anthropocene. Science, 345(6195), 401–406. https://doi.org/10.1126/science.1251817

Duffy, J. E., Godwin, C. M., & Cardinale, B. J. (2017). Biodiversity effects in the wild are common and as strong as key drivers of productivity. Nature, 549(7671), 261–264. https://doi.org/10.1038/nature23886

Eigenbrod, F., Hecnar, S. J., & Fahrig, L. (2011). Sub-optimal study design has major impacts on landscape-scale inference. Biological Conservation, 144(1), 298–305. https://doi.org/10.1016/j.biocon.2010.09.007

Fahrig, L., Baudry, J., Brotons, L., Burel, F. G., Crist, T. O., Fuller, R. J., Sirami, C., Siriwardena, G. M., & Martin, J.-L. (2011). Functional landscape heterogeneity and animal biodiversity in agricultural landscapes. Ecology Letters, 14(2), 101–112. https://doi.org/10.1111/j.1461-0248.2010.01559.x

Foley, J. A., DeFries, R., Asner, G. P., Barford, C., Bonan, G., Carpenter, S. R., Chapin, F. S., Coe, M. T., Daily, G. C., Gibbs, H. K., Helkowski, J. H., Holloway, T., Howard, E. A., Kucharik, C. J., Monfreda, C., Patz, J. A., Prentice, I. C., Ramankutty, N., & Snyder, P. K. (2005). Global consequences of land use. Science, 309(5734), 570–574. https://doi.org/10.1126/science.1111772

Fox, R., Oliver, T. H., Harrower, C., Parsons, M. S., Thomas, C. D., & Roy, D. B. (2014). Long-term changes to the frequency of occurrence of British moths are consistent with opposing and synergistic effects of climate and land-use changes. Journal of Applied Ecology, 51(4), 949–957. https://doi.org/10.1111/1365-2664.12256

Gillespie, M. A. K., Baude, M., Biesmeijer, J., Boatman, N., Budge, G. E., Crowe, A., Memmott, J., Morton, R. D., Pietravalle, S., Potts, S. G., Senapathi, D., Smart, S. M., & Kunin, W. E. (2017). A method for the objective selection of landscape-scale study regions and sites at the national level. Methods in Ecology and Evolution, 8(11), 1468–1476. https://doi.org/10.1111/2041-210X.12779

Hallmann, C. A., Sorg, M., Jongejans, E., Siepel, H., Hofland, N., Schwan, H., Stenmans, W., Müller, A., Sumser, H., Hörren, T., Goulson, D., & Kroon, H. de. (2017). More than 75 percent decline over 27 years in total flying insect biomass in protected areas. PLOS ONE, 12(10), e0185809. https://doi.org/10.1371/journal.pone.0185809

Harrison, S. (2020). Plant community diversity will decline more than increase under climatic warming. Philosophical Transactions of the Royal Society B: Biological Sciences, 375(1794), 20190106. https://doi.org/10.1098/rstb.2019.0106

Holzschuh, A., Dainese, M., González-Varo, J. P., Mudri-Stojnić, S., Riedinger, V., Rundlöf, M., Scheper, J., Wickens, J. B., Wickens, V. J., Bommarco, R., Kleijn, D., Potts, S. G., Roberts, S. P. M., Smith, H. G., Vilà, M., Vujić, A., & Steffan-Dewenter, I. (2016). Mass-flowering crops dilute pollinator abundance in agricultural landscapes across Europe. Ecology Letters, 19(10), 1228–1236. https://doi.org/10.1111/ele.12657

Jochner, S., Caffarra, A., & Menzel, A. (2013). Can spatial data substitute temporal data in phenological modelling? A survey using birch flowering. Tree Physiology, 33(12), 1256–1268. https://doi.org/10.1093/treephys/tpt079

Laux, P., Nguyen, P. N. B., Cullmann, J., Van, T. P., & Kunstmann, H. (2017). How many RCM ensemble members provide confidence in the impact of land-use land cover change? International Journal of Climatology, 37(4), 2080–2100. https://doi.org/10.1002/joc.4836

Martin, E. A., Dainese, M., Clough, Y., Báldi, A., Bommarco, R., Gagic, V., Garratt, M. P. D., Holzschuh, A., Kleijn, D., Kovács-Hostyánszki, A., Marini, L., Potts, S. G., Smith, H. G., Hassan, D. A., Albrecht, M., Andersson, G. K. S., Asís, J. D., Aviron, S., Balzan, M. V., … Steffan-Dewenter, I. (2019). The interplay of landscape composition and configuration: New pathways to manage functional biodiversity and agroecosystem services across Europe. Ecology Letters, 22(7), 1083–1094. https://doi.org/10.1111/ele.13265

Mori, A. S., Isbell, F., & Seidl, R. (2018). β-Diversity, community assembly, and ecosystem functioning. Trends in Ecology & Evolution, 33(7), 549–564. https://doi.org/10.1016/j.tree.2018.04.012

Newbold, T., Hudson, L. N., Hill, S. L. L., Contu, S., Lysenko, I., Senior, R. A., Börger, L., Bennett, D. J., Choimes, A., Collen, B., Day, J., De Palma, A., Díaz, S., Echeverria-Londoño, S., Edgar, M. J., Feldman, A., Garon, M., Harrison, M. L. K., Alhusseini, T., … Purvis, A. (2015). Global effects of land use on local terrestrial biodiversity. Nature, 520(7545), 45–50. https://doi.org/10.1038/nature14324

Parmesan, C. (2006). Ecological and Evolutionary Responses to Recent Climate Change. Annual Review of Ecology, Evolution, and Systematics, 37(1), 637–669. https://doi.org/10.1146/annurev.ecolsys.37.091305.110100

Pasher, J., Mitchell, S. W., King, D. J., Fahrig, L., Smith, A. C., & Lindsay, K. E. (2013). Optimizing landscape selection for estimating relative effects of landscape variables on ecological responses. Landscape Ecology, 28(3), 371–383. https://doi.org/10.1007/s10980-013-9852-6

Peters, M. K., Hemp, A., Appelhans, T., Becker, J. N., Behler, C., Classen, A., Detsch, F., Ensslin, A., Ferger, S. W., Frederiksen, S. B., Gebert, F., Gerschlauer, F., Gütlein, A., Helbig-Bonitz, M., Hemp, C., Kindeketa, W. J., Kühnel, A., Mayr, A. V., Mwangomo, E., … Steffan-Dewenter, I. (2019). Climate–land-use interactions shape tropical mountain biodiversity and ecosystem functions. Nature, 568(7750), 88–92. https://doi.org/10.1038/s41586-019-1048-z

Piano, E., Souffreau, C., Merckx, T., Baardsen, L. F., Backeljau, T., Bonte, D., Brans, K. I., Cours, M., Dahirel, M., Debortoli, N., Decaestecker, E., Wolf, K. D., Engelen, J. M. T., Fontaneto, D., Gianuca, A. T., Govaert, L., Hanashiro, F. T. T., Higuti, J., Lens, L., … Hendrickx, F. (2020). Urbanization drives cross-taxon declines in abundance and diversity at multiple spatial scales. Global Change Biology, 26(3), 1196–1211. https://doi.org/10.1111/gcb.14934

Provost, G. L., Badenhausser, I., Bagousse-Pinguet, Y. L., Clough, Y., Henckel, L., Violle, C., Bretagnolle, V., Roncoroni, M., Manning, P., & Gross, N. (2020). Land-use history impacts functional diversity across multiple trophic groups. Proceedings of the National Academy of Sciences, 117(3), 1573–1579. https://doi.org/10.1073/pnas.1910023117

Rabe, S.-E., Koellner, T., Marzelli, S., Schumacher, P., & Grêt-Regamey, A. (2016). National ecosystem services mapping at multiple scales—The German exemplar. Ecological Indicators, 70, 357–372. https://doi.org/10.1016/j.ecolind.2016.05.043

Redlich, S., Martin, E. A., & Steffan-Dewenter, I. (2018). Landscape-level crop diversity benefits biological pest control. Journal of Applied Ecology, 55(5), 2419–2428. https://doi.org/10.1111/1365-2664.13126

Rillig, M. C., Ryo, M., Lehmann, A., Aguilar-Trigueros, C. A., Buchert, S., Wulf, A., Iwasaki, A., Roy, J., & Yang, G. (2019). The role of multiple global change factors in driving soil functions and microbial biodiversity. Science, 366(6467), 886–890. https://doi.org/10.1126/science.aay2832

Seibold, S., Gossner, M. M., Simons, N. K., Blüthgen, N., Müller, J., Ambarlı, D., Ammer, C., Bauhus, J., Fischer, M., Habel, J. C., Linsenmair, K. E., Nauss, T., Penone, C., Prati, D., Schall, P., Schulze, E.-D., Vogt, J., Wöllauer, S., & Weisser, W. W. (2019). Arthropod decline in grasslands and forests is associated with landscape-level drivers. Nature, 574(7780), 671–674. https://doi.org/10.1038/s41586-019-1684-3

SRTM. (2020). Digital Elevation - Shuttle Radar Topography Mission (SRTM) 1 Arc-Second. https://doi.org/10.5066/TEEBF7PR7TFT

TEEB. (2010). The economics of ecosystems and biodiversity. Ecological and economic foundations. edited by Pushpam Kumar. Earthscan London, Washington

Thies, C., Steffan-Dewenter, I., & Tscharntke, T. (2003). Effects of landscape context on herbivory and parasitism at different spatial scales. Oikos, 101(1), 18–25. https://doi.org/10.1034/j.1600-0706.2003.12567.x

Urban, M. C. (2015). Accelerating extinction risk from climate change. Science, 348(6234), 571–573. https://doi.org/10.1126/science.aaa4984

Wagner, D. L. (2020). Insect declines in the Anthropocene. Annual Review of Entomology, 65(1), 457–480. https://doi.org/10.1146/annurev-ento-011019-025151

Warren, M. S., Hill, J. K., Thomas, J. A., Asher, J., Fox, R., Huntley, B., Roy, D. B., Telfer, M. G., Jeffcoate, S., Harding, P., Jeffcoate, G., Willis, S. G., Greatorex-Davies, J. N., Moss, D., & Thomas, C. D. (2001). Rapid responses of British butterflies to opposing forces of climate and habitat change. Nature, 414(6859), 65–69. https://doi.org/10.1038/35102054

Warren, R., Price, J., Graham, E., Forstenhaeusler, N., & VanDerWal, J. (2018). The projected effect on insects, vertebrates, and plants of limiting global warming to 1.5°C rather than 2°C. Science, 360(6390), 791–795. https://doi.org/10.1126/science.aar3646

Wiens, J. A. (1989). Spatial scaling in ecology. Functional Ecology, 3(4), 385–397. https://doi.org/10.2307/2389612

Williams, J. J., & Newbold, T. (2020). Local climatic changes affect biodiversity responses to land use: A review. Diversity and Distributions, 26(1), 76–92. https://doi.org/10.1111/ddi.12999

