## Appendix for "Disentangling effects of climate and land use on biodiversity and ecosystem services – a multi-scale experimental design"

Redlich et al. 2021

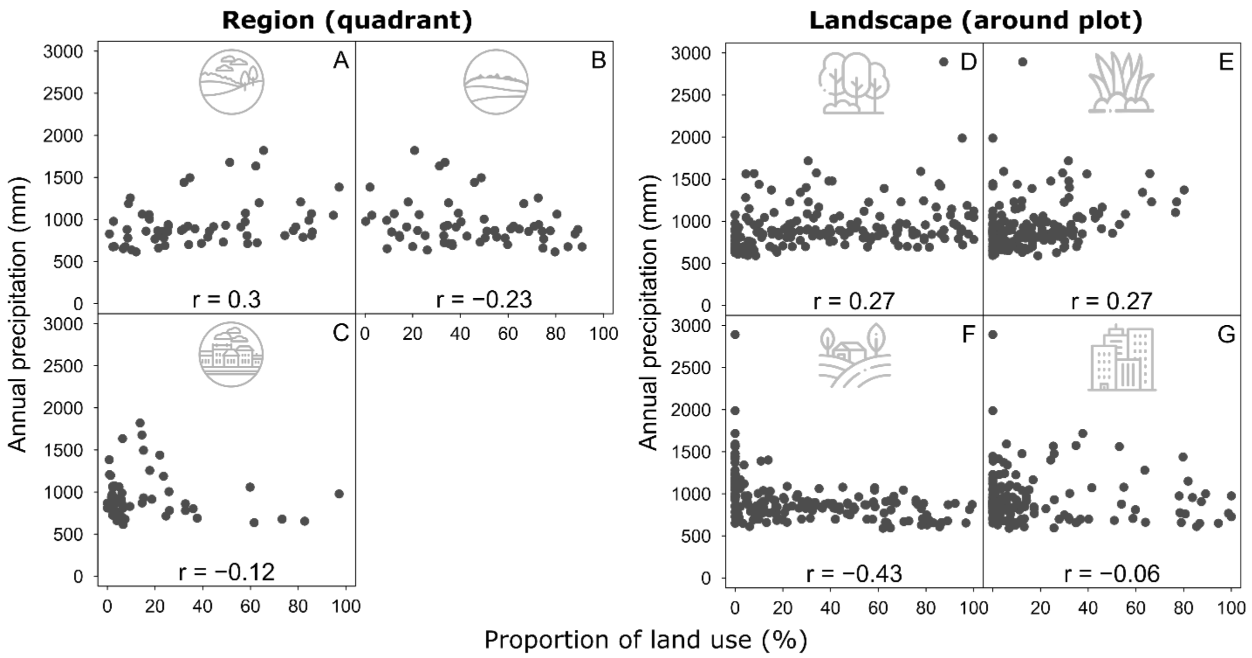


Figure S1. Relationships between annual precipitation (mm) and proportional land cover (%) for the regional land-use types near-nature (A), agriculture (B) and urban (C), and for the landscape-scale land-use types forest (D), grassland (E), arable land (F) and settlement (G). Pearson's *r* coefficients based on 60 study regions (5.8 x 5.8 km quadrants, A–C) and 179 study plots (1 km radius around local study plots, D–G).

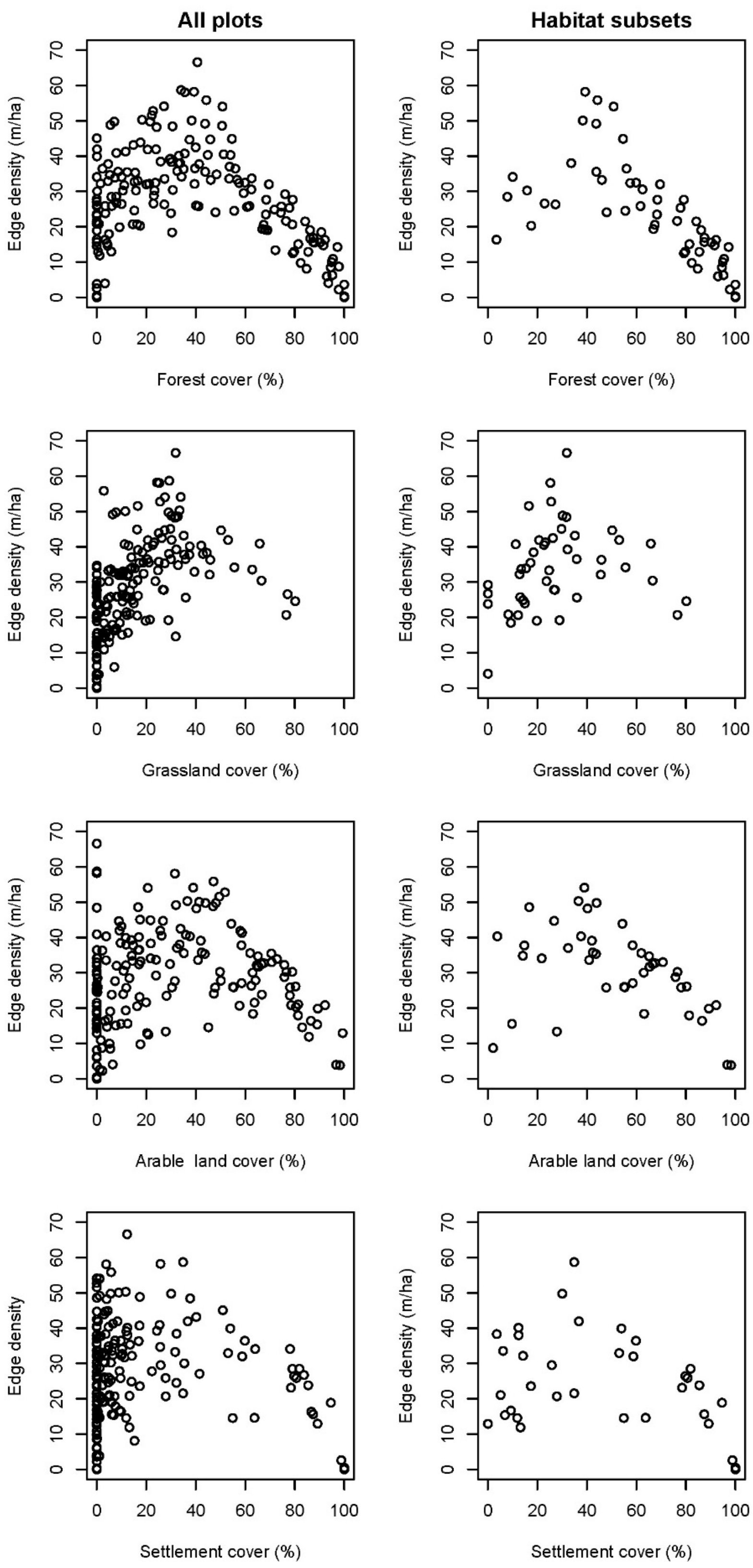

All plots
Habitat subsets
Edge density (m/ha)
Forest cover (%)
Grassland cover (%)
Arable land cover (%)
Edge density
Settlement cover (%)
0
10
20
30
40
50
60
70
0
20
40
60
80
100

Figure S2. Relationships between landscape composition and edge density across all 179 study plots and only considering plots of associated local habitat type (e.g. correlation between total edge density and forest composition for forest plots only).
